# Murine cytomegalovirus *m119-118* encodes a new viral Fc receptor antagonizing host Fc receptor activation

**DOI:** 10.64898/2026.10.09.758151

**Authors:** Laura Staliunaite, Maria Mellin, Eléonore Ostermann, Katrin Ehrhardt, Baca Chan, Alec J. Redwood, Andrew J. Davison, Felix R. Stahl, Philipp Kolb, Hartmut Hengel, Wolfram Brune

**Affiliations:** Leibniz Institute of Virology (LIV), 20251 Hamburg, Germany; Institute of Virology, Medical Center and Faculty of Medicine, University of Freiburg, 79104 Freiburg, Germany; School of Biomedical Science, University of Western Australia, Nedlands, WA 6009, Australia; Institute for Respiratory Health, University of Western Australia, Nedlands, WA 6009, Australia; MRC-University of Glasgow Centre for Virus Research, Glasgow G61 1QH, UK; Institute of Immunology, University Medical Center Hamburg-Eppendorf, 20251 Hamburg, Germany; Hamburg Center for Translational Immunology, University Medical Center Hamburg-Eppendorf, 20251 Hamburg, Germany

## Abstract

Cytomegaloviruses (CMVs) encode numerous immunomodulatory proteins to counteract innate and adaptive immunity. One class of immune evasion proteins is viral Fc receptors (vFcRs), which bind to the Fc region of IgG antibodies and interfere with antibody-mediated effector functions crucial for antiviral defense. Despite the established role of vFcRs in modulating antibody-mediated immunity, their contribution to viral pathogenesis in vivo remains incompletely understood. Although human CMV (HCMV) encodes four vFcRs, only a single vFcR, fcr-1 encoded by gene *m138*, has been identified in murine CMV (MCMV). By comparing and cloning the genomes of MCMV strains v70, K181^+^, and K181^Perth^, we identified a frameshift mutation affecting the spliced *m119-118* transcript in K181^Perth^. This finding prompted us to investigate the function of the *m119-118* gene. We show that it encodes a second MCMV vFcR, which we named fcr-2. It is structurally similar to its positional homolog in HCMV, UL119-118/gp68, and binds specifically to the Fc part of murine IgG. Deletion of both vFcRs genes from the MCMV genome resulted in substantially reduced Fc-binding to the surface of infected cells, indicating a complementary function of the vFcRs. Functional assays with viral deletion mutants further demonstrated that both proteins inhibit host Fc receptor activation in a cooperative fashion. Although an m119-118-deficient MCMV was only marginally attenuated in infected BALB/c mice, an m119-118 and m138 double mutant attained significantly lower organ titers than the m138-deficient mutant. These data suggest an important cooperative function of the two MCMV vFcRs for viral dissemination and pathogenesis in vivo.

**Author Summary:** Cytomegaloviruses (CMVs) establish lifelong infections by evading elimination by the host immune system. One strategy used to achieve this is the expression of proteins that bind antibodies and interfere with antibody-mediated immune responses. Although several antibody-binding proteins have been identified in human CMV, only one has been described in mouse cytomegalovirus (MCMV), m138/fcr-1. In this study, we identified a second antibody-binding protein encoded by gene *m119-118*, which we name fcr-2. We show that this protein binds IgG antibodies and acts together with m138/fcr-1 to suppress activation of host antibody-binding receptors. In mice, deletion of *m119-118* alone had only a minor effect on viral infection. In contrast, deletion of both genes caused substantially greater defects in replication and spread than viruses lacking either gene alone. These findings demonstrate that MCMV uses at least two cooperating antibody-binding proteins to weaken antibody-dependent immune defenses and promote infection in vivo.

## Introduction

Human cytomegalovirus (HCMV) is a widespread pathogen that establishes lifelong latent infection in humans and can cause severe disease in immunocompromised individuals and congenitally infected infants [1]. Approved antiviral treatment options remain limited, and no vaccine is currently available [2]. Despite the robust humoral and cellular immune responses induced during primary infection, reinfection with antigenically distinct HCMV strains occurs frequently throughout life, underscoring the virus’s remarkable ability to evade pre-existing immunity [3,4]. To establish lifelong persistence and facilitate repeated infections, HCMV has evolved a broad array of immune evasion mechanisms that interfere with the activation and effector functions of both innate and adaptive immune responses [5,6].

Although potent neutralizing antibodies are regularly present in the sera of previously infected individuals, the clinical efficacy of antibody-based interventions against HCMV has remained limited [7]. This suggests that HCMV efficiently evades or counteracts antibody-mediated immune functions beyond virus neutralization, such as antibody-dependent cellular phagocytosis (ADCP) and cytotoxicity (ADCC) [8,9]. Four HCMV genes encode viral Fc receptors (vFcRs): *UL119-118* (encoding glycoprotein gp68), *RL11* (encoding glycoprotein gp34), *RL12*, and *RL13* [10–12]. These receptors have been shown to bind to the Fc portion of human IgG, thereby interfering with the activation of host Fc receptors (FcRs) expressed on immune cells such as NK cells, macrophages, monocytes, and dendritic cells [13,14]. For gp68 and gp34, immune antagonization has been shown to involve distinct mechanisms, including antibody bipolar bridging (ABB), which sterically impairs FcR engagement, as well as the internalization of immune complexes from the cell surface. By preventing FcR activation, the vFcRs impair downstream effector mechanisms such as ADCC [13,14].

Experimental studies of HCMV infection and pathogenesis in vivo are impracticable as HCMV is a human-specific virus that does not infect other animals [15]. For this reason, the role of HCMV vFcRs in viral dissemination and pathogenesis in vivo has remained poorly understood. However, recent studies with the related rhesus CMV (RhCMV) have identified three vFcRs encoded by *Rh05*, *Rh152-151*, and *Rh173* genes [16,17]. Infection experiments in immunocompetent, RhCMV-naïve rhesus macaques showed that deleting single vFcR genes from the RhCMV genome did not reduce viral fitness in vivo, but deleting all three vFcRs significantly shortened DNAemia [17].

Infection of mice with murine CMV (MCMV) is a well-established and practical model for studying CMV pathogenesis in vivo [18]. To date, only one MCMV-encoded vFcR has been identified: it is encoded by the *m138* gene, which is also known as *fcr-1* [19]. Deletion of *m138* from the MCMV genome resulted in a marked attenuation of viral dissemination in vivo [20,21]. However, subsequent studies revealed that the m138 protein fulfills several additional immune evasion functions: it downregulates the NKG2D ligands MULT-1, RAE-1, and H60 to dampen NK cell activation [20,22,23]. It also downregulates costimulatory molecules B7-1 and ICOSL to reduce T-cell activation [24–26]. These antibody-independent functions are predominantly responsible for the marked attenuation of the *m138*-deleted MCMV in vivo [21]. These and other observations suggested that antibody-dependent immune responses may play only a minor role during primary MCMV infection [27]. Alternatively, MCMV might encode additional vFcRs that partially compensate for one another when individual Fc-binding proteins are absent, as it has been shown in HCMV and RhCMV [14,17].

Strains Smith and K181 are the best characterized and most widely used MCMV strains [28–30]. K181 was first isolated in the laboratory of June Osborn [29]. Plaque-purified K181 isolates have been described, such as K181^+^ [31] and K181^Perth^ [32]. The genome of K181^Perth^ has also been cloned as a bacterial artificial chromosome (BAC) in *E. coli* [33]. Another isolate, called v70, also originated in the Osborn lab. It has been described as a more virulent variant of the Smith strain [34–36]. However, genetic studies to define its ancestry have not yet been done.

In this study, we sequenced the genomes of MCMV strains K181^+^, K181^Perth^, and v70, and cloned them as BACs using the STAR cloning method [37,38]. We show that these strains are very similar and represent variants of strain K181. Sequence comparisons revealed that K181^Perth^ contains a frameshift mutation in the previously uncharacterized gene *m119-118*. Using structural and functional analyses, we show that *m119-118* encodes a novel vFcR that acts together with m138/fcr-1 to repress host FcR activation in vitro and contributes to viral fitness in vivo.

## Results

### Genetic comparison and STAR cloning of MCMV strains K181^+^, K181^Perth^, and v70

MCMV strain v70 was generated by serial propagation through the salivary glands of CD-1 mice [34]. It has been used in numerous immunological studies, but its ancestry has remained unclear, particularly since it was repeatedly described as a variant of the Smith strain, even though it originated from the same laboratory that isolated the K181 strain. To resolve this enigma, we determined the genome sequence of the v70 virus by Illumina sequencing. For comparison, we sequenced the genomes of K181^+^ and K181^Perth^ isolates. An alignment of the three viral genome sequences revealed that v70 is genetically highly similar to both K181 isolates. Compared to K181^+^, v70 differs by only three nucleotides: one T-tract variation in a noncoding region located between M105 and M112, and two single nucleotide polymorphisms (SNPs) leading to silent mutations in M43 and M105, respectively (Table 1). The genome sequence of the K181^Perth^ isolate differed from the K181^Perth^ reference sequence (GenBank accession AM886412) only by a single three-nucleotide deletion in the *m131* gene, leading to the loss of a glycine residue (ΔGly95) in the MCK-2 protein. The K181^Perth^ and v70 sequences differ in five positions (Table 1). Three differences caused SNPs or nucleotide tract variations, and two differences were considered of greater importance: in addition to the *m131* deletion mentioned above, K181^Perth^ contains a single-nucleotide deletion in *m119*, leading to a frameshift and early truncation of the protein encoded by the spliced *m119-118* gene (Table 1). In contrast, the v70 genome differs in more than 1500 positions from the MCMV strain Smith genome. Hence, we concluded that v70 is a variant of the K181 strain, just like K181^+^ and K181^Perth^.

**Table 1.**
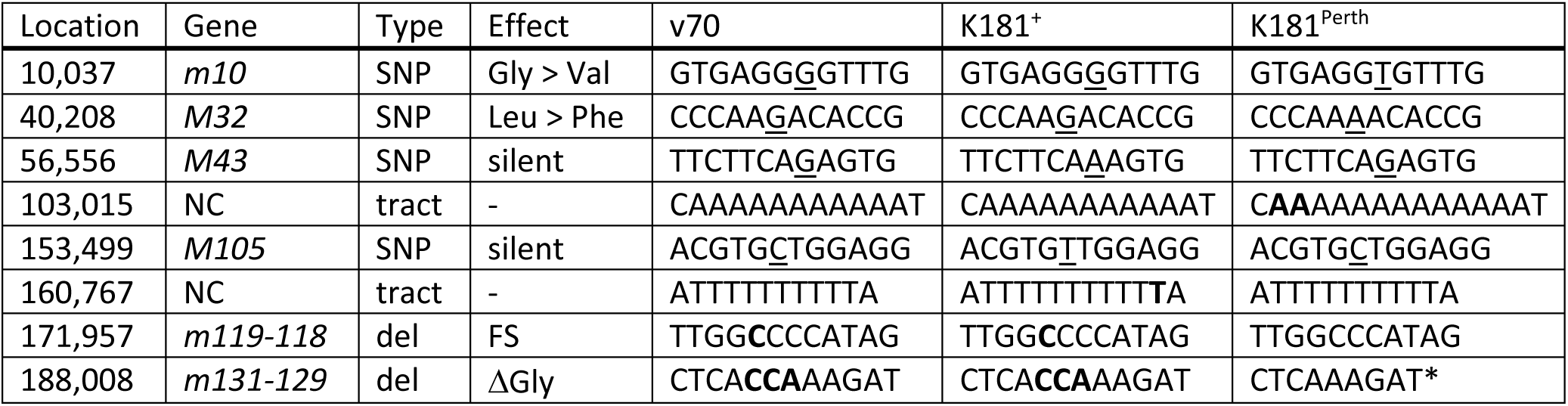
Genetic differences between MCMV strains v70, K181+, and K181^Perth^. Nucleotide positions (Location) refer to the v70 genome (GenBank accession QB061181). NC, non-coding region; SNP, single-nucleotide polymorphism; del, deletion; FS, frameshift; ΔGly, loss of glycine residue due to three-nucleotide deletion. Amino acid substitutions are indicated by >. SNPs are underlined, and tract variations and deletions are in bold font; *, deletion present in the K181^Perth^ isolate used in this study, but not in the K181^Perth^ reference sequence (GenBank AM886412).

To investigate the functional relevance of strain-specific mutations and variations, we made the three K181-derived strains amenable to genetic modification. To do this, we generated BAC clones of v70, K181^+^, and K181^Perth^ using single-step transformation-associated recombination (STAR) cloning, which was recently developed in our lab [37,38]. The pCC1BAC-his3 cloning vector was modified by inserting 60-bp hooks homologous to the ends of the viral genome, as done previously for the cloning of the rat CMV and *Mastomys natalensis* CMV genomes [37,39]. Full-length viral genomes isolated from viral particles and the linearized cloning vector were used for transformation of *S. cerevisiae*, where transformation-associated recombination took place (Fig. 1A). Yeast clones harboring the cloned MCMV genome were identified by PCR screening. PCR-positive BACs were isolated from yeast and transferred to *E. coli*. The completeness and integrity of the BAC-cloned viral genomes were first analyzed by restriction digest and gel electrophoresis (Fig. 1B) and subsequently verified by complete genome sequencing. The K181^Perth^ BAC sequence was identical to the consensus sequence of the parental virus, but the K181^+^ and v70 BAC sequences differed from the parental virus genomes in 3 and 1 positions, respectively. These deviations were repaired by *en passant* BAC mutagenesis [40]. The integrity of the repaired BAC clones was verified by complete BAC sequencing. Infectious virus was reconstituted in 10.1 fibroblasts and used for virus stock production.

**Figure 1.**
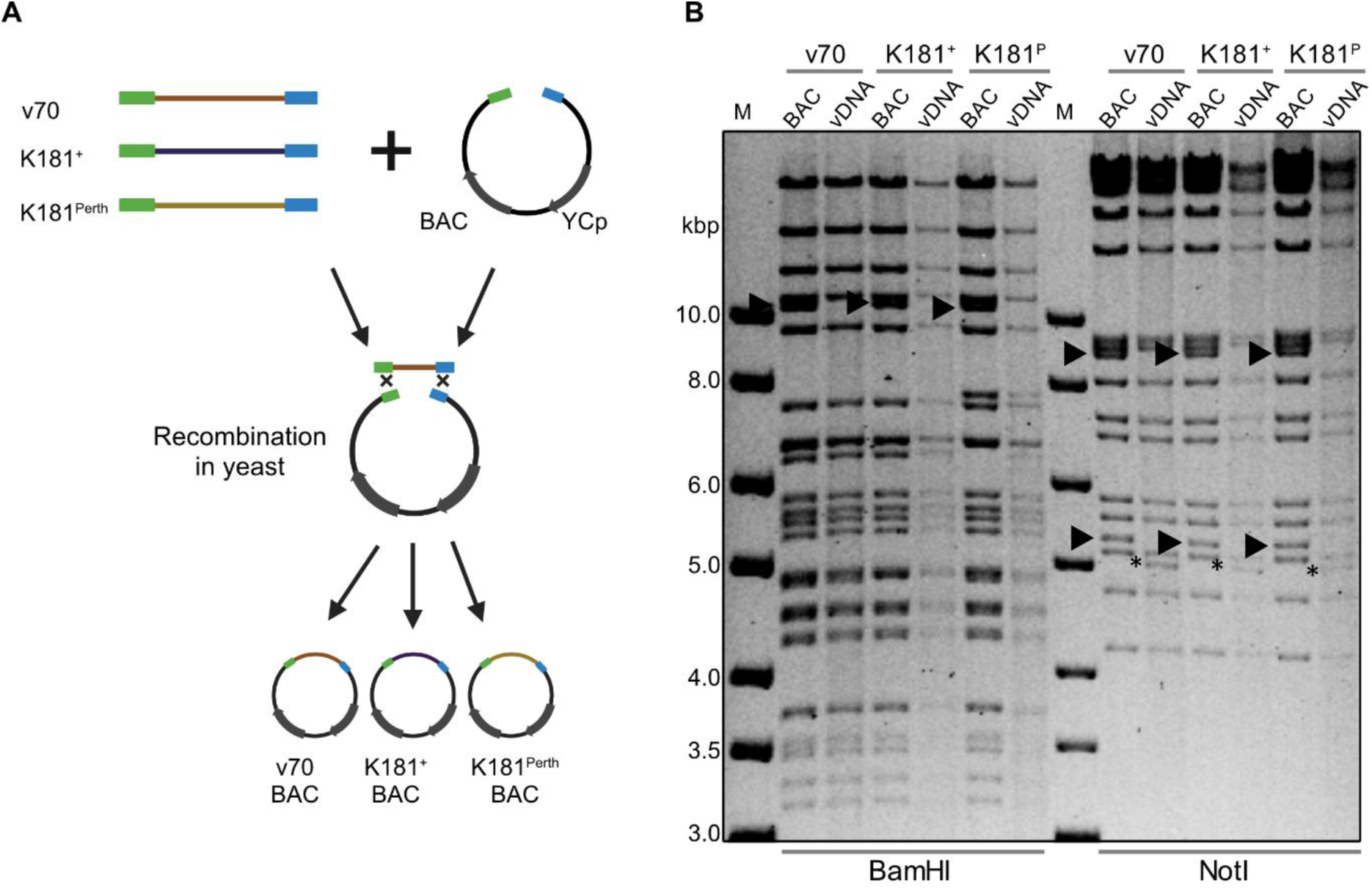
STAR cloning of MCMV strains. (**A**) Linear MCMV DNA isolated from virions (vDNA) and linearized cloning vector were used for homologous recombination in yeast. Homology hooks (60 bp) corresponding to the viral genome ends are shown in green and blue. Circular plasmids consisting of the vector and the full-length MCMV genome were isolated from yeast. Schematic created with BioRender. (**B**) BAC DNA of cloned MCMV genomes and vDNA used for cloning were digested with BamHI or NotI and analyzed by gel electrophoresis. Vector-vDNA junction fragments (arrowheads) and vDNA terminal fragments (asterisks) are indicated. M, molecular weight marker.

### The ΔGly95 mutation in MCK-2 of K181^Perth^ does not affect viral replication in macrophages

The v70 strain has preserved its biological fitness by serial propagation in mice, whereas other strains may have acquired mutations during propagation in cell culture. Therefore, we tested whether mutations identified by sequence comparisons (Table 1) have an impact on viral fitness. First, we focused on a 3-bp deletion in the K181^Perth^ *m131* gene, which leads to the loss of a glycine residue (ΔGly95) in the MCK-2 protein. MCK-2 forms a trimeric glycoprotein complex with glycoprotein H (gH) and glycoprotein L (gL), which is important for the infection of macrophages in vitro and in vivo and promotes viral dissemination to the salivary glands [41–45]. Thus, we assessed viral infectivity and replication of MCMV v70, K181^+^, and K181^Perth^ in J774A.1 macrophages, immortalized bone marrow derived macrophages (iBMDMs), and 10.1 fibroblasts (as a control). However, no significant differences in infectivity and viral replication in macrophages were detected (S1 Fig), suggesting that the loss of a single glycine residue in MCK-2 did not impair the ability of K181^Perth^ to infect macrophages.

### MCMV *m119-118* encodes a glycoprotein that interacts with the Fc part of mouse IgG

Next, we focused on the frameshift mutation in K181^Perth^ *m119*. This ORF was reported to be part of a spliced *m119-118* transcript [46,47], but its protein product has not been analyzed. The *m119-118* gene is a positional homolog of the HCMV *UL119-118* gene, but amino acid similarity between the two proteins is low (23% identity; 35% similarity). However, the structures predicted by AlphaFold3 were quite similar, consisting of an α-helical segment and two β-sheet rich regions each [48], (Fig. 2A). As *UL119-118* encodes a glycoprotein (gp68) that functions as a vFcR and antagonizes host FcR activation [13,14,49], we tested whether the m119-118 protein also encodes a vFcR. To do this, we cloned the *m119-118* gene with a C-terminal HA tag in a plasmid vector (pcDNA3). As controls, we also cloned *m138* (encoding the only known MCMV vFcR) and the glycoprotein B (gB) gene with a C-terminal FLAG tag in the same vector. After transient transfection of HEK293 cells with these expression plasmids, cell lysates were incubated with mouse IgG. Immunoprecipitation with protein G sepharose and immunoblot analysis showed that m119-118 and m138 bind to mouse IgG whereas gB does not (Fig. 2B).

**Figure 2.**
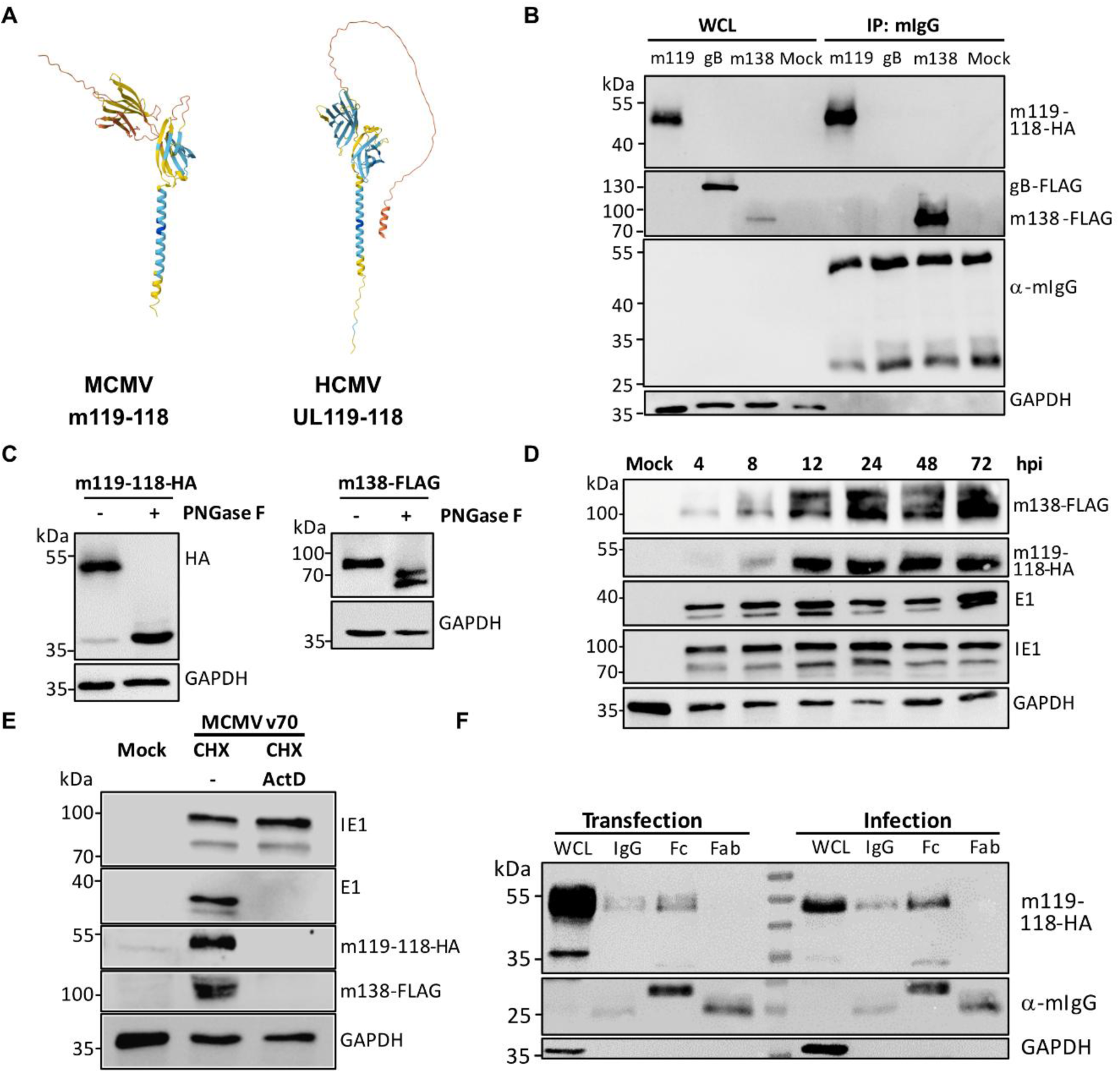
Expression kinetics and Fc binding of m119-118 and m138. **(A)** Structure predictions of MCMV m119-118 and HCMV UL119-118/gp68 proteins generated using AlphaFold3. Structures are colored according to the pLDDT (predicted Local Distance Difference Test) score, with blue indicating high confidence and red indicating low confidence. **(B)** HEK293 cells were transfected with plasmids expressing m119-118-HA, m138-FLAG, or gB-FLAG. Whole-cell lysates were subjected to immunoprecipitation with mouse IgG. Precipitated proteins were detected by Western blot analysis. Antibodies against GAPDH and mouse IgG served as loading and precipitation controls, respectively. **(C)** HEK293 cells were transfected with the same plasmids, and lysates were digested with PNGase F. Treated and untreated samples were analyzed by Western blot. **(D)** Murine 10.1 fibroblasts were infected with MCMV v70 m119-118-HA/m138-FLAG (MOI=1), and lysates were harvested at various times post-infection and subjected to SDS-PAGE. Antibodies against GAPDH, IE1, and E1 served as loading and infection controls, respectively. **(E)** 10.1 fibroblasts infected with MCMV m119-118-HA/138-FLAG were treated for 4 hours with 50 μg/mL cycloheximide (CHX) followed by 5 μg/mL Actinomycin D (ActD) or no (–) treatment for 5 hours prior to Western blot analysis. **(F)** HEK293 cells were transfected with an m119-118-HA plasmid, and 10.1 murine fibroblasts were infected with MCMV v70 m119-118-HA (MOI=1) for 48 h. Whole cell lysates were subjected to immunoprecipitation with total mouse IgG or biotinylated Fc or Fab fragments. Precipitated proteins were detected by Western blot analysis.

As the apparent molecular weight of the m119-118 protein (approx. 50 kDa) did not match the predicted molecular weight of 33 kDa, cell lysates were treated with PNGase F to remove glycosylation. After treatment, the apparent molecular weight was reduced to approximately 35 kDa, which is close to the predicted weight (Fig. 2C). A molecular weight reduction upon PNGase F treatment was also confirmed for m138 (Fig. 2C), indicating that both proteins are glycosylated.

Next, we analyzed the function of m119-118 in the context of viral infection. Using *en passant* BAC mutagenesis, we modified the v70 BAC by inserting an HA tag sequence at the 3’ end of *m119-118* and a FLAG tag sequence at the 3’ end of *m138*. The kinetics of m119-118 and m138 protein expression in fibroblasts infected with the tagged virus was analyzed by Western blot (Fig. 2D). The detection of both proteins as early as 4-8 hours post-infection (hpi) suggested that the genes are expressed with early kinetics. By using a cycloheximide release assay, we showed that both proteins are not expressed at immediate-early times (Fig. 2E).

The tagged v70 virus was also used to test whether the m119-118 protein expressed in MCMV-infected cells binds to IgG and whether it binds to the Fab or the Fc part. A pull-down experiment was done with lysates of plasmid-transfected cells and MCMV-infected cells. In both cases, m119-118 bound to IgG and specifically to the Fc but not the Fab fragment (Fig. 2F). These results suggested that the *m119-118* gene encodes a vFcR, which we propose to call fcr-2.

### The two vFcRs, fcr-1 and fcr-2, synergistically inhibit host FcR activation

To further analyze the importance of the two vFcRs, we generated a set of v70 deletion mutants by BAC mutagenesis. Flow cytometric analysis and viral replication kinetics in 10.1 fibroblasts showed that the single (Δm119-118 and Δm138) and double deletion (ΔΔm119/138) mutants showed similar infectivity compared to WT and were not replication-impaired (Fig. 3A, B). These mutants were then used to measure Fc binding to the surface of infected fibroblasts. Cells were infected with v70 WT, Δm119-118, Δm138, or ΔΔm119/138 for 24 h and incubated with biotinylated Fc fragments. After incubation with PE-labeled streptavidin, Fc surface binding was assessed by flow cytometry (Fig. 3C, D). The result showed that Fc binding was slightly reduced in cells infected with Δm119-118 and more significantly reduced in cells infected with Δm138. The loss of Fc binding was most pronounced in cells infected with ΔΔm119/m138, reducing mean fluorescence intensity (MFI) by approximately 75%. This indicates that both vFcRs are present on the surface of infected cells and can functionally bind IgG Fc fragments. However, Fc binding was not completely abolished in cells infected with the ΔΔm119/m138 mutant, as residual binding remained greater than mock levels.

**Figure 3.**
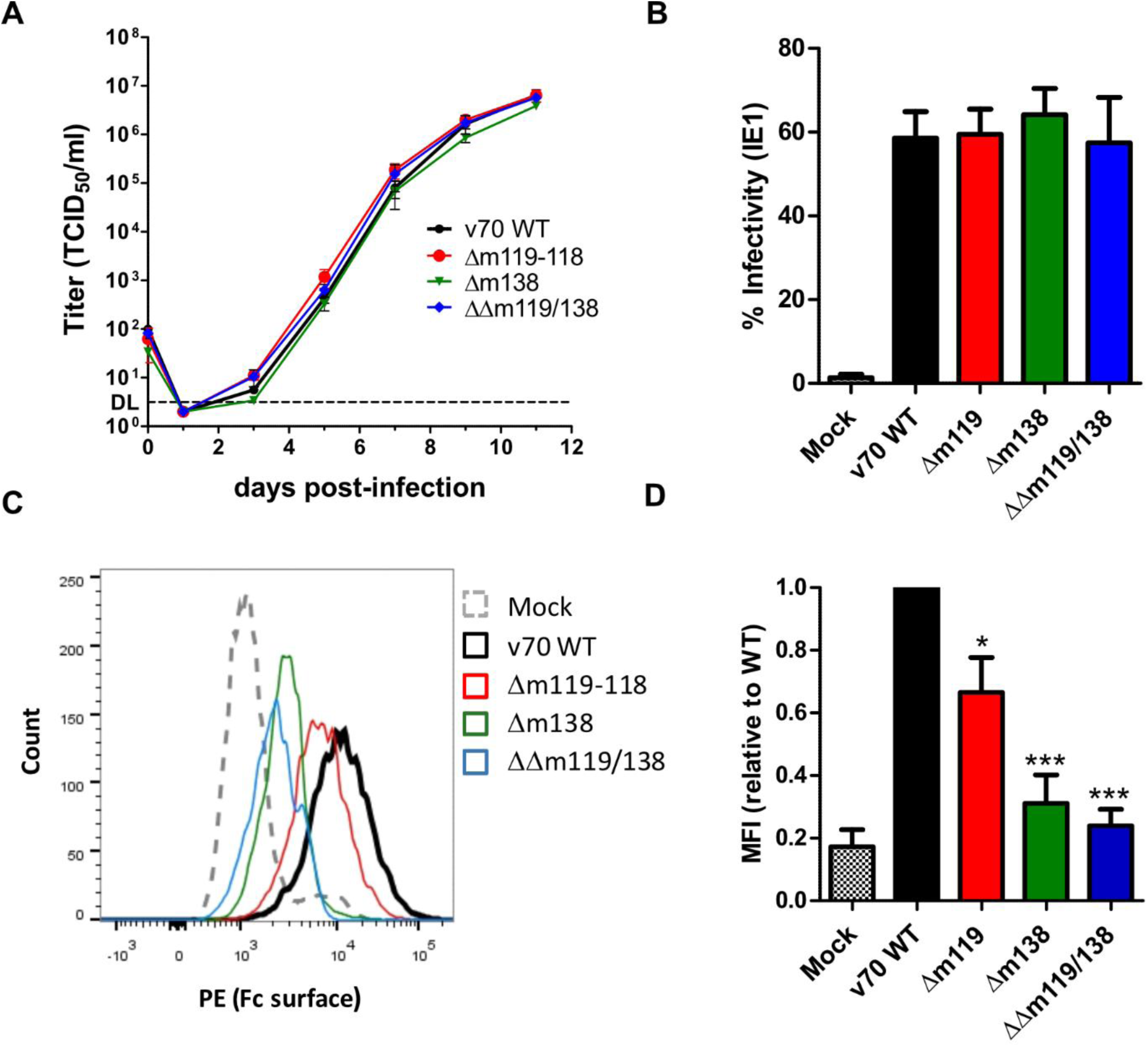
Replication, infectivity, and Fc surface binding of vFcR mutants. **(A)** 10.1 fibroblasts were infected with BAC-derived MCMV v70 WT or mutants (MOI=0.02). Viral titers in the supernatants were determined using the TCID_50_ method. Mean ± SEM of three biological replicates is shown. DL, detection limit. **(B)** 10.1 fibroblasts were infected with WT v70 or the respective mutants for 24 h (MOI=1), fixed, permeabilized, and stained with an IE1-specific antibody. Infectivity was assessed by flow cytometry. Mean ± SEM of three individual experiments is shown. **(C)** 10.1 fibroblasts were infected as described in (B). Cells were incubated with biotinylated Fc fragments and phycoerythrin (PE)-coupled streptavidin. Fc surface binding was analyzed and quantified by flow cytometry. One of three independent experiments is shown. Mean ± SEM of all three experiments is shown in **(D)** as mean fluorescence intensity (MFI), relative to WT. Significance was calculated using one-way ANOVA with Bonferroni’s multiple comparison test. Asterisks indicate adjusted *p* values. *, *p* < 0.05; ***, *p* < 0.001.

To assess whether the vFcRs expressed by MCMV-infected cells can antagonize host FcR activation, we used a previously developed FcR reporter assay [50,51]. Modified BW5147 lymphoblast lines expressing mouse FcRI (CD64), FcRIII (CD16), FcRIV, or the inhibitory receptor FcRIIB (CD32), fused to the transmembrane and cytoplasmatic domains of mouse CD3ζ, were used for this assay (Fig. 4A). In this assay, binding of IgG-containing immune complexes to chimeric host FcRs induces receptor activation, phosphorylation of the CD3ζ domain and subsequent interleukin 2 (IL-2) secretion, which can be quantified as a measure of FcR activation. We therefore used this assay to determine whether the MCMV vFcRs can compete with host FcRs for IgG binding and thereby reduce host FcR activation.

**Figure 4.**
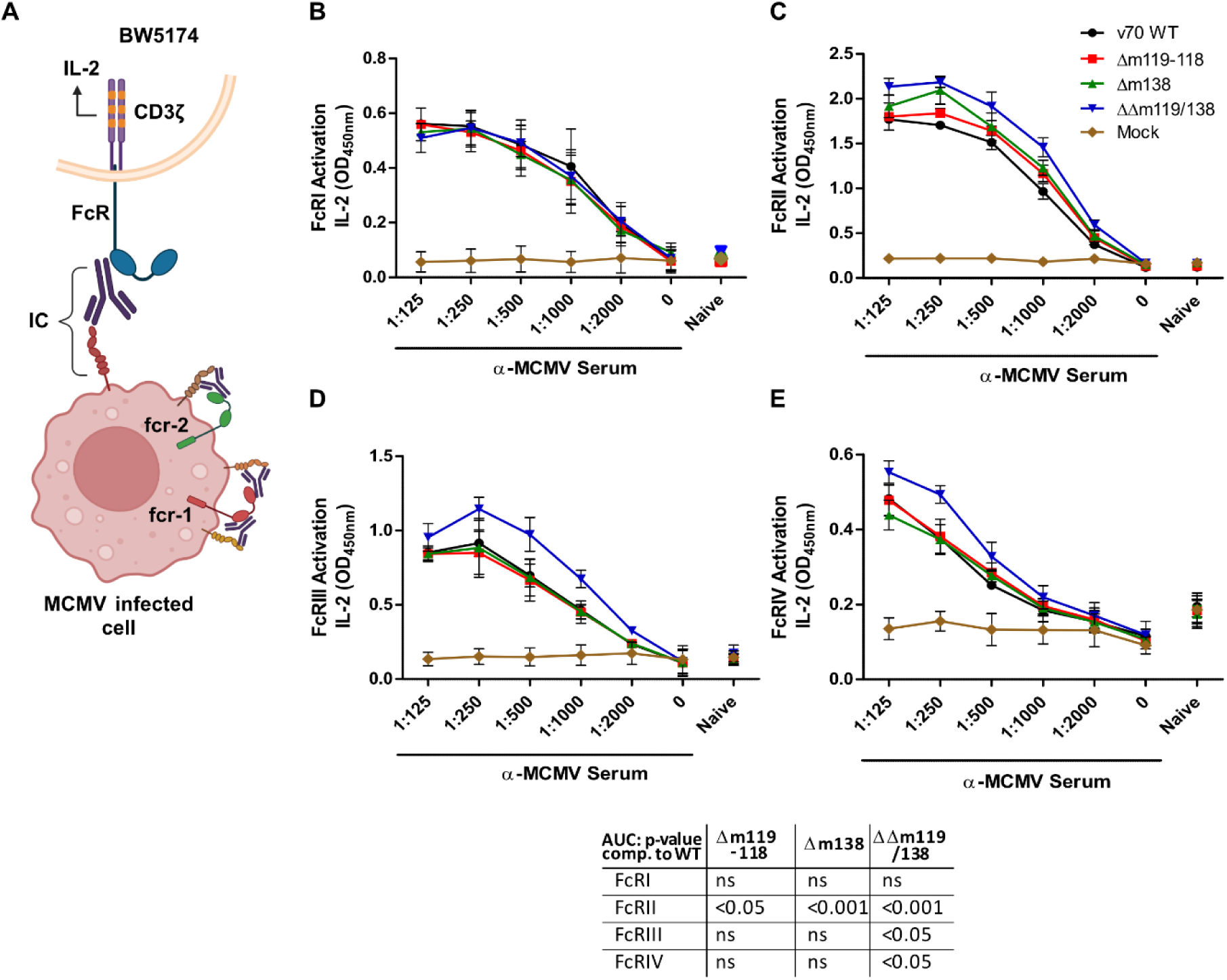
Antagonization of host FcR activation by MCMV vFcRs expressed in infected fibroblasts. **(A)** Schematic of BW5147 reporter cell assay. A mouse FcR extracellular domain is fused to the transmembrane and intracellular domains of CD3ζ, secreting IL-2 upon activation. The receptor is activated by interaction to immune complexes (ICs). MCMV infected cells express vFcRs, binding antibodies via the Fc fragment and counteracting host FcR activation. Created with BioRender. (**B-E**) 10.1 fibroblasts were infected with WT or mutant v70 strains (MOI=1) for 24 h. Uninfected cells served as control. Anti-MCMV antiserum was added in various dilutions. Serum from uninfected mice served as a negative control. BW5147 reporter cells ectopically expressing host FcRs I (B), II (C), III (D) or IV (E) were added, and activation of host FcRs was measured 18 h later using an IL-2 ELISA. Mean ±SEM of 2 or 3 independent experiments is shown. For statistical analysis, the area under curve (AUC) was calculated for each dataset. The *p* values shown in the table were determined by one-way ANOVA and adjusted by Bonferroni’s multiple correction. ns, not significant.

First, the impact of vFcRs expressed by MCMV-infected cells on host FcR activation was analyzed. Fibroblasts were infected with MCMV v70 WT or the deletion mutants and incubated with anti-MCMV antiserum (from previously infected BALB/c mice) for immune complex formation. Mock-infected cells and serum from naïve animals served as controls. Host FcR-expressing BW5147 cells were added, and the activation of host FcRs was analyzed by measuring IL-2 secretion. Compared to WT v70-infected fibroblasts, activation of FcRII, FcRIII, and FcRIV was significantly increased when fibroblasts were infected with the ΔΔm119/138 mutant (Fig. 4B-E). Deletion of only one vFcR had less or no effect on host FcR activation, suggesting that the two vFcRs have additive and partly redundant functions. Interestingly, activation of host FcRI was not affected by the presence or absence of the two vFcRs in infected fibroblasts.

### Viral FcRs incorporated in viral particles inhibit host FcR activation

Next, we wanted to find out whether the vFcRs are incorporated into viral particles (virions) and whether virion-incorporated vFcRs can inhibit FcR activation. Virions of MCMV expressing tagged vFcRs (see Fig. 2D) were purified using discontinuous Nycodenz gradient centrifugation, lysed, and analyzed by Western Blot (Fig. 5A). A strong m119-118-specific band was detected in virion lysates, suggesting that substantial quantities of fcr-2 were present in viral particles. The m138/fcr-1 protein was also detected in purified virions, but the signal was weak in repeated experiments, suggesting that the protein is incorporated into virions at relatively low levels. The purity of the virion preparation was assessed by detecting GAPDH and IE1 (present in whole cell lysates but not in virions). As expected, purified virions contained only cleaved gB (Fig. 5A).

**Figure 5.**
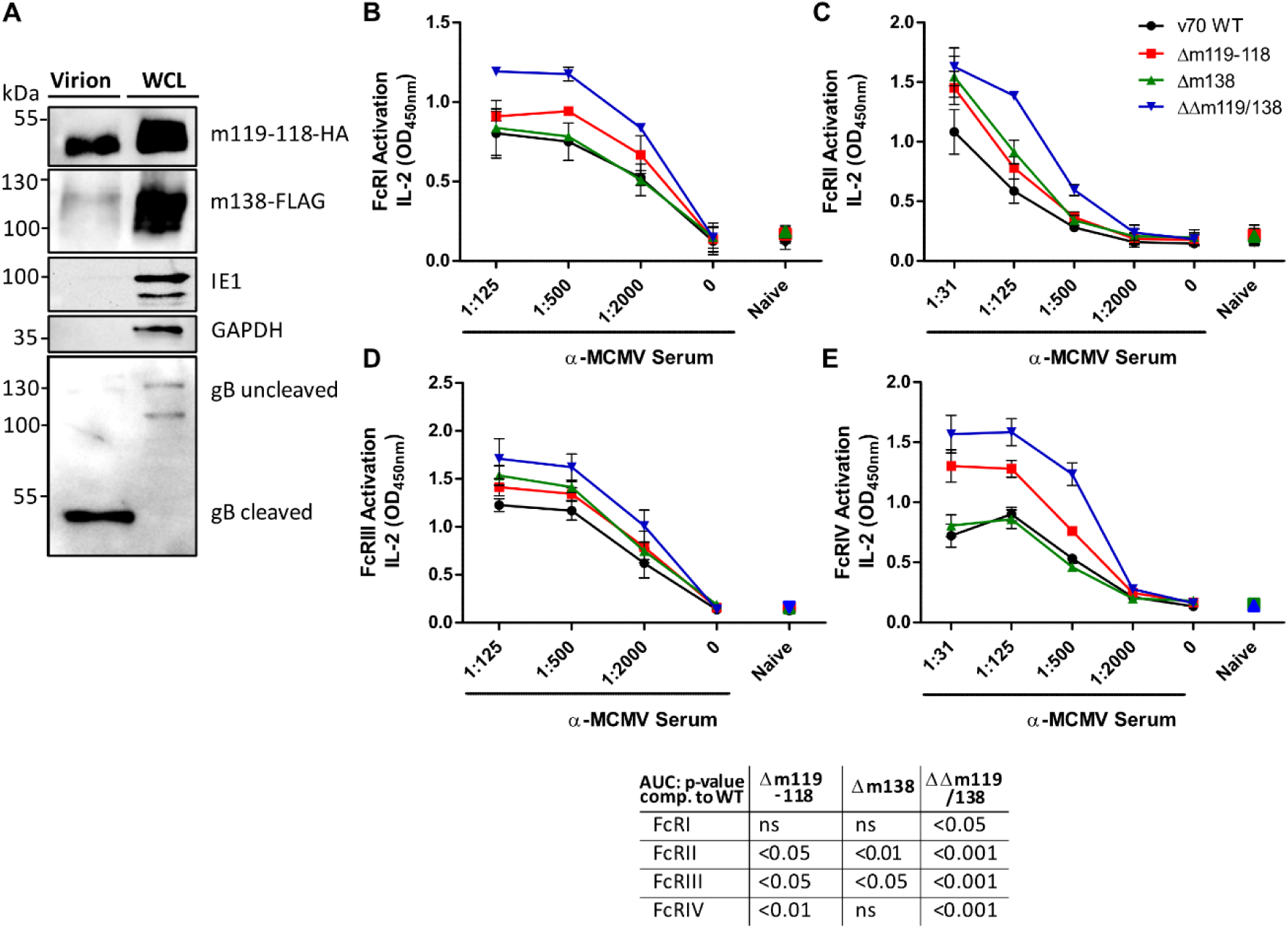
MCMV vFcRs in virions inhibit host FcR activation. (**A**) Viral particles of MCMV v70 m119-118-HA/m138-FLAG purified by Nycodenz gradient centrifugation. Purified virions and whole cell lysates (WCL) were analyzed by Western blot. (**B-E**) Equal amounts of virion protein was used for ELISA plate coating. Anti-MCMV antiserum was added in various dilutions. Serum from uninfected mice was used as a negative control. BW5147 reporter cells were added, and activation of host FcRs was measured 18 h later using an IL-2 ELISA. Mean ±SEM of 2 or 3 independent experiments is shown. For statistical analysis, the area under curve (AUC) was calculated for each dataset. The *p* values shown in the table were determined by one-way ANOVA and adjusted by Bonferroni’s multiple correction. ns, not significant.

To test whether vFcRs in viral particles can inhibit host FcR activation, we used the same FcR assay as above but coated microtiter plates with viral particles instead of MCMV-infected cells. To ensure that equivalent amounts of viral particles were used for each virus, viral protein content was quantified with an anti-MCMV ELISA (S2 Fig.). Virion-coated microtiter plates were incubated with anti-MCMV antiserum, and FcR-expressing BW5147 cells were added. Host FcR activation was analyzed by measuring IL-2 secretion. The activation of all four host FcRs was significantly increased when ΔΔm119/138 virions were used (Fig. 5B-E). The increase was lower or insignificant with Δm119-118 or Δm138 virions.

### Viral FcRs inhibit host FcR activation by antibodies against a non-viral target

To find out whether MCMV vFcRs specifically antagonize host FcR activation by MCMV-specific antibodies, we used an antibody recognizing a non-viral target. Murine fibroblasts stably expressing human CD8 were infected with v70 WT and mutant MCMVs, incubated with a mouse anti-hCD8 IgG antibody, and co-cultured with BW5147 Fc reporter cells (Fig. 6A-D). Again, host FcR activation was significantly increased when cells were infected with the mutant lacking both vFcR genes. The effect was lower when cells were infected with single vFcR-deficient mutants. Hence, we concluded that the inhibition of host FcR activation by vFcRs is not restricted to MCMV-specific antibodies.

**Figure 6.**
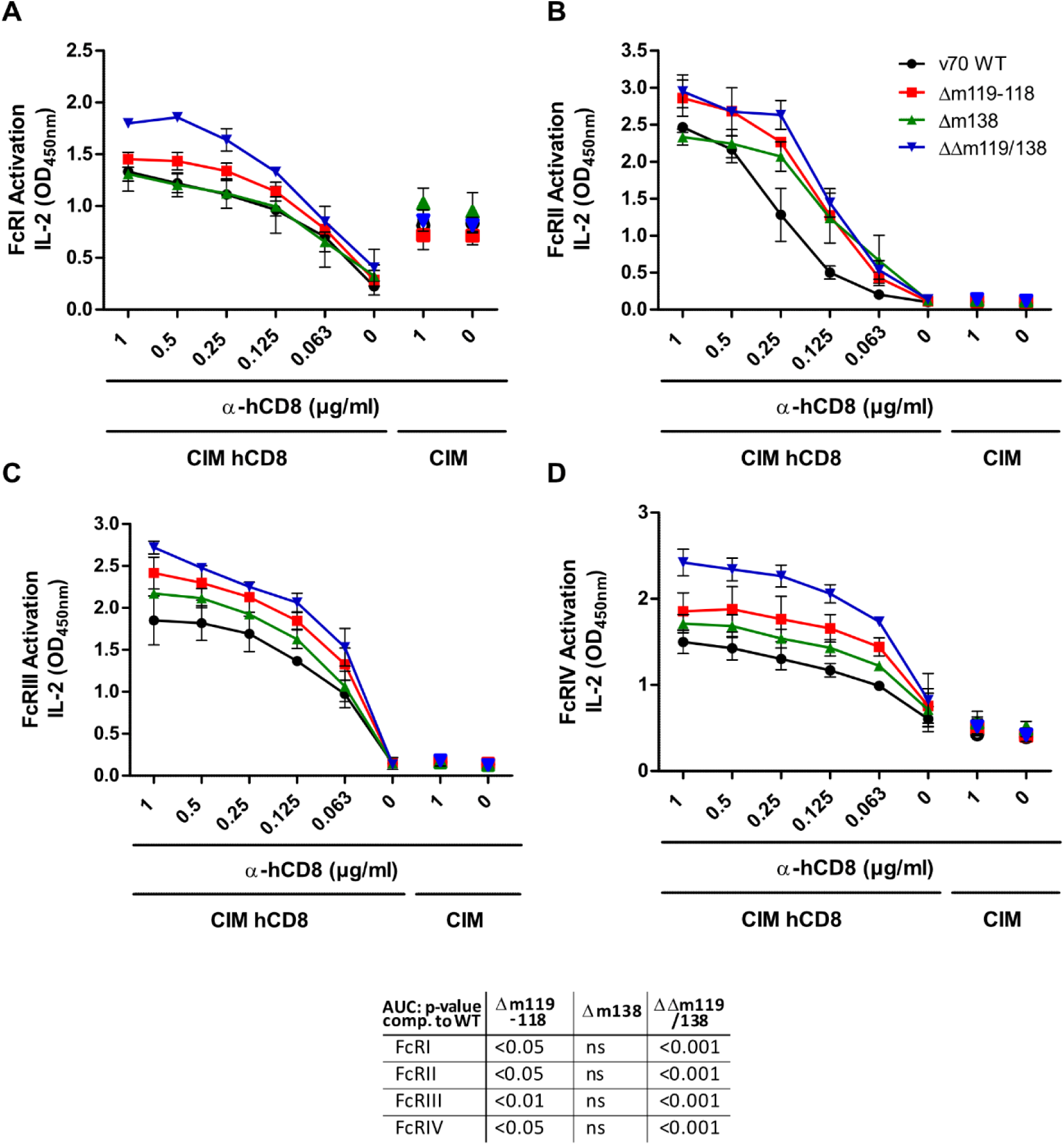
MCMV vFcRs antagonize host FcR activation by MCMV-unrelated immune complexes. (**A-D**) WT and hCD8-expressing CIM cells were infected with WT and mutant MCMV v70 strains (MOI=1) for 18 h. Murine anti-hCD8 antibodies were added in various concentrations. BW5147 reporter cells were added, and host FcR activation was measured 18 h later using an IL-2 ELISA. Mean ± SEM of 3 independent experiments is shown. For statistical analysis, the area under curve (AUC) was calculated for each dataset. The *p* values shown in the table were determined by one-way ANOVA and adjusted by Bonferroni’s multiple correction. ns, not significant.

### Viral FcRs are required for viral fitness in vivo

To determine the roles of the two vFcRs, m119-118 and m138, in MCMV replication and dissemination in mice, a set of mutant MCMVs suitable for in vivo experiments was needed. The mutants used in the previous experiments carried deletions of the entire ORFs that might affect the expression of neighboring genes. In contrast, less disruptive mutations are less likely to have polar effects. We introduced stop mutations into ORFs *m119* and *m138* after codons 58 and 56, respectively. The stop mutations were first introduced into v70 BAC expressing tagged vFcRs by *en passant* BAC mutagenesis. As an additional control, the *m119* frameshift mutation of K181^Perth^ was introduced. Fibroblasts were infected with the recombinant viruses, and vFcR expression was analyzed by Western blot. As expected, the stop mutations completely abolished the expression of the respective proteins. The same loss of fcr-2 expression was achieved by the frameshift mutation (Fig. 7A). We concluded that the stop mutations were suitable for in vivo experiments and introduced the same mutations into the v70 BAC to obtain m119*stop*, m138*stop*, and m119/138*stop* mutants. The mutant viral genomes were sequenced completely to exclude the presence of adventitious mutations elsewhere in the viral genome.

**Figure 7.**
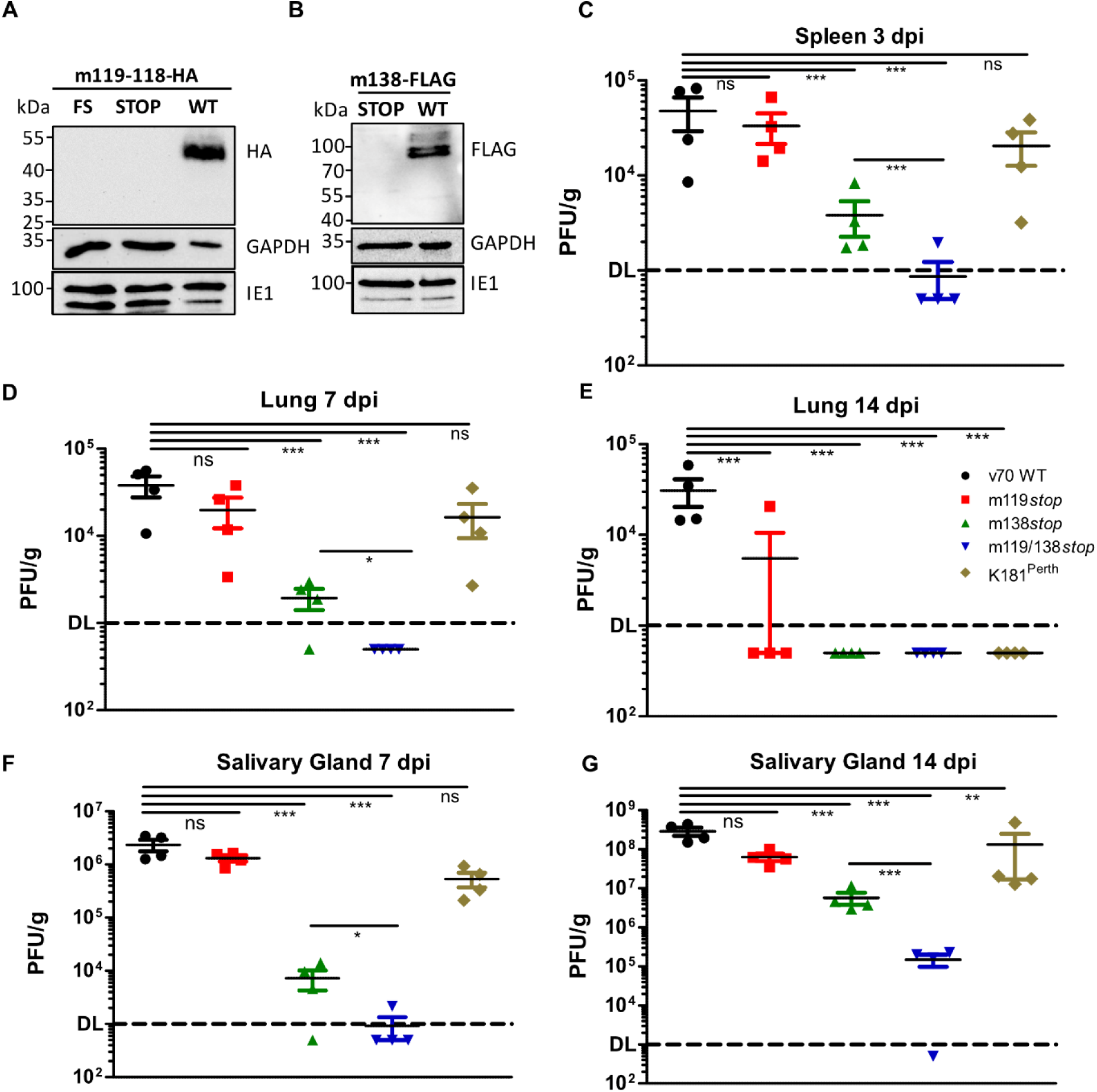
MCMV replication and dissemination *in vivo* in the presence or absence of vFcRs. (**A**) 10.1 fibroblasts were infected (MOI=1) with MCMV v70 expressing C-terminally HA-tagged m119-118 (WT), or mutants additionally carrying either an early stop codon or the K181^Perth^-specific frameshift (FS) mutation in *m119-118*. Cells lysates were harvested 36 hpi and analyzed by Western blot. IE1 and GAPDH were included as infection and loading controls, respectively. (**B**) 10.1 fibroblasts were infected with MCMV expressing C-terminally FLAG-tagged m138 (WT) or the corresponding stop mutant. Infection and Western blot analysis was done as described above. (**C-G**) BALB/c mice were infected i.p. with 2×10^5^ PFU of WT or mutant MCMV v70. Mice were sacrificed on days 3, 7, and 14 post-infection, and viral titers in organs were determined by plaque assay. Each symbol represents an individual animal (n=4 per group), bars indicate mean ± SEM. Statistical significance was calculated using two-way ANOVA and Bonferroni’s multiple comparison of log-transformed grouped data. Asterisks indicate adjusted *p* values: *, *p* < 0.05; **, *p* < 0.01; ***, *p* < 0.001. ns, not significant.

BALB/c mice were infected intraperitoneally with WT and mutant MCMV v70 strains and with K181^Perth^, which contains a frameshift mutation in *m119* gene. Mice were sacrificed on days 3, 7, and 14 post-infection, and viral titers in the spleen, lungs, and salivary glands were determined. Organ titers of the m119*stop* mutant were slightly reduced compared to WT v70, but the reduction was statistically significant only in the lung at 14 dpi (Fig. 7B-E). A similar small titer reduction was detected after infection with K181^Perth^. As expected, the loss of m138 alone led to a severe attenuation with 10-to 100-fold reduced titers compared to WT v70 in all organs. This strong attenuation is consistent with previous reports and largely depends on additional antibody-independent functions of m138 [20,21,26]. Importantly, inactivation of both vFcR genes (m119/m138*stop*) caused a significantly stronger attenuation than inactivation of only one vFcR gene (Fig. 6B-E). In line with results obtained with host FcR activation assays (Figs. 3, 4, and 5), these findings suggest a cooperative function of the two vFcRs to support viral replication and dissemination *in vivo*.

## Discussion

In this study, we report the characterization of MCMV m119-118/fcr-2 as a novel vFcR that, together with the previously identified vFcR m138/fcr-1, inhibits host FcR activation. Host FcR activation assays with MCMV mutants lacking *fcr-1*, *fcr-2*, or both genes revealed that deletion of one vFcR gene only moderately increased host FcR activation, whereas the deletion of both genes resulted in a more pronounced increase. It is noteworthy that the contribution of m138/fcr-1 was stronger when host FcR activation was induced by antibodies bound to MCMV-infected cells (Fig. 4). Conversely, the inhibitory effect of m119-118/fcr-2 was stronger when activation was induced by antibodies bound to viral particles (Fig. 5). The high abundance of the fcr-2 in viral particles is a plausible explanation for this disparity. Interestingly, neither fcr-2 nor fcr-1 was detected in a mass spectrometric analysis of the MCMV virion proteome [52]. However, the HCMV vFcR gp68, encoded by *UL119-118*, was detected in virions by both protein mass spectrometry and Western blot analysis [53,54]. MCMV *m119-118* is a positional homolog of HCMV *UL119-118*, and the two proteins exhibit remarkably similar Fc-binding activities despite their limited sequence similarity. Consequently, m119-118 provides a useful model for investigating the in vivo function of UL119-118/gp68. Such studies cannot be performed directly with HCMV, as the virus is strictly human-specific and does not infect animals. Recently, the RhCMV *Rh152-151* gene was also characterized as a functional vFcR and ortholog of *UL119-118* [17], demonstrating that FcR homologues are conserved across multiple CMVs. Although RhCMV infection of rhesus macaques may offer the closest approximation to HCMV infection in vivo, particularly in the context of congenital CMV infection, its use is limited by ethical considerations, high costs, limited animal availability, and the lack of genetically modified models. MCMV infection of mice will therefore remain the most practical, accessible, and experimentally tractable system for investigating the role of vFcRs in CMV pathogenesis.

The fact that HCMV and RhCMV express four and three vFcRs, respectively [14,17] raises the question of whether MCMV also encodes additional vFcRs besides m138/fcr-1 and m119-118/fcr-2. Indeed, Fc binding activity, as measured by flow cytometry, was significantly reduced but not completely abolished in cells infected with the ΔΔm119/138 mutant (Fig. 3C, D). The residual Fc binding may indicate the presence of an additional MCMV vFcR. Alternatively, it may also be caused by technical limitations of the assay or increased nonspecific staining of MCMV-infected cells. Deletion of all four HCMV encoded vFcRs however reduced Fc surface binding completely to mock levels [14].

Until now, the contribution of MCMV vFcRs to viral pathogenesis could only be studied to a limited extent because m138/fcr-1 was the only known Fc-binding protein. Since fcr-1 also downregulates NK- and T-cell ligands [20,23,24,26], the strong attenuation of m138 deletion mutants cannot be attributed solely to the loss of FcR modulation [21]. Thus, the identification of fcr-2 provides a new opportunity to investigate the importance of viral FcRs for pathogenesis in vivo. In fact, inactivation of fcr-2 alone resulted in only minor attenuation. However, a pronounced reduction in viral fitness was observed when fcr-2 was inactivated in addition to fcr-1 (Fig. 7). These results suggested that the two vFcRs have additive and partially redundant functions in vivo, similar to their functional activity in the host FcR activation assays (Figs. 4-6).

The best way to address the question of how Fc-binding by viral proteins contributes to pathogenesis in vivo is to introduce subtle mutations into the vFcRs that abrogate Fc binding without affecting other functions of the viral proteins. This is a challenging task as the various functions may require the same protein domains. For instance, Fc binding and downregulation of MULT-1, H60, B7-1, and ICOSL depend on the N-terminal Ig1 domain of m138 [20,24,26]. Once a specifically Fc binding-deficient mutant becomes available, any remaining attenuation can be attributed to an antibody-mediated mechanism, particularly if it is rescued in antibody-deficient mice.

Another interesting question to be addressed in future studies is the apparent attenuation of the m119/138*stop* mutant in the spleen at 3 dpi. As mice start to produce virus-specific IgG antibodies only 5-7 days after MCMV infection [55], this attenuation cannot be attributed to MCMV-specific antibodies. However, it is possible that such early effects are mediated by natural antibodies (NAbs) or natural autoantibodies (NAAbs) [56,57]. Whereas NAbs are low-affinity antibodies that (cross-)react with viral antigens, NAAbs may recognize stress antigens presented by virus-infected cells. However, little is known about the contribution of NAbs and NAAbs to the control of MCMV infection thus far. Alternatively, the attenuation at very early times might be caused by hitherto unknown additional functions of m119-118/fcr-2.

The identification of fcr-2 as a functional MCMV vFcR also highlights the importance of strain-specific differences in virus research. Sequencing of strains v70, K181^+^, and K181^Perth^ revealed a high degree of genetic similarity, suggesting a very close relationship between the strains. However, sequencing also identified a previously unrecognized frameshift mutation in *m119-118* in K181^Perth^, which results in the complete loss of fcr-2 protein expression. Notably, comparative analysis of various published MCMV genome sequences revealed that fcr-2 is highly conserved across other MCMV genomes, including the Smith, G4, WP15, HaNa1, and SN1 strains, showing at least 94% amino acid sequence identity. This high degree of conservation suggests that fcr-2 may be under functional constraint and could indicate an important role of the vFcR across different MCMV strains. As K181^Perth^ and its BAC clone, pARK25, are widely used laboratory strains, the identified mutation should be considered, particularly for in vivo studies. Although deletion of *fcr-2* results in only a subtle phenotype in an otherwise wild-type virus, its combination with additional mutations may result in a markedly enhanced phenotype, as observed in the present study.

HCMV is known to be genetically unstable when propagated in cell culture. It rapidly acquires mutations, particularly in RL13 and the UL128-131 regions [58]. In contrast, MCMV has been considered relatively stable during passage in cell culture. This notion may have to be revised. In several instances, frameshift mutations have been identified in cell culture-propagated MCMV strains: a frameshift mutation in *m129-131*/*mck-2* was identified in the MCMV Smith strain [41], a frameshift mutation in *m133-132*/*sgg1.1* was found in the G4 strain [59], and a frameshift mutation in *m119-118*/*fcr-2* in the K181^Perth^ strain was described in the present study.

This study identified the MCMV m119-118/fcr-2 as a new vFcR, characterized its activity as an antagonist of host FcR activation, and revealed its importance for MCMV infection and dissemination *in vivo*. First studies on previously uncharacterized viral genes cannot answer all questions, and this is also true for the present study. One important question is whether the pronounced attenuation of the ΔΔm119/138 mutant, compared to the m138 single deletion mutant, depends on the IgG-binding activities of the two viral vFcRs or on additional antibody-independent functions. For m138/fcr-1, several antibody-independent functions have been described [20,23,24,26], and these additional functions are largely responsible for the strong attenuation of m138-deficient MCMV mutants [21]. Whether the glycoprotein encoded by *m119-118* also has antibody-independent functions is currently unknown. The minimal attenuation of the m119*stop* mutant *in vivo* (Fig. 6) suggests that there are no additional functions of major importance for viral fitness. Yet, additional functions cannot be completely excluded. So far, no Fc-independent functions have been described for HCMV UL119-118/gp68 or RhCMV Rh152-151 [14,17]. However, it remains possible that such additional functions exist and become apparent only under specific infection conditions in vivo. As in vivo experiments are impossible for HCMV and severely limited for RhCMV, the identification of m119-118/*fcr-2* as the second MCMV-encoded vFcR and ortholog of HCMV UL119-118 and RhCMV Rh152-151 opens new opportunities to investigate the biological importance of vFcRs in vivo, particularly when combined with the genetically modified mouse models available for infection research.

## Material and Methods

### Cells and viruses

M2-10B4 bone marrow stromal cells (CRL-1972) and HEK293 cells (CL-11268) were obtained from the American Type Culture Collection. Murine J774A.1 macrophages (ECACC 91051511) were obtained from the European Collection of Authenticated Cell Cultures, and immortalized bone marrow-derived macrophages (iBMDM, NR-9456) from BEI Resources. 10.1 fibroblasts are spontaneously immortalized MEFs from BALB/c mice [60]. CIM cells are mouse fibroblasts isolated from C57BL/6 embryonic tissue and were kindly provided by Dr. Mirko Trilling (University of Duisburg-Essen, Germany). Cells were cultured at 37°C and 5% CO_2_ in Dulbecco’s modified Eagle medium (DMEM), containing 10% FCS, 100 IU/mL penicillin, and 100 µg/mL streptomycin.

CIM cells expressing hCD8 were generated by lentiviral transduction and puromycin selection as described [61]. CD8 expression was verified by flow cytometry using an APC-labeled anti-CD8 antibody (OKT-8, eBioscience, S3 Fig.). Murine BW5147 lymphoblast cells (ATCC TIB-47) stably expressing mouse FcRs have been described previously [50,51]. They were grown in RPMI 1640 GlutaMAX medium, supplemented with 10% FCS, 50 IU/mL penicillin, 50 µg/ml streptomycin, 1% sodium pyruvate, and 100 mM β-mercaptoethanol.

MCMV strain v70 was serially passaged in mice by Dr. Maryjane Selgrade (Environmental Protection Agency, Research Triangle Park, NC)[34] and provided to Dr. Christine Biron (Brown University, Providence, RI)[62]. Dr. Biron provided the v70 strain to Dr. Edward Mocarski (Emory University, Atlanta, GA)[36], who kindly provided it to us. The K181^Perth^ strain was also provided by Dr. Edward Mocarski. K181^+^ is a 3× plaque-purified isolate of the MCMV K181 strain [31], which was provided to us by Dr. Jason Upton (Auburn University, Auburn, AL). MCMV was grown on 10.1 fibroblasts, and virus stocks were prepared following standard protocols [63].

### Plasmids

The following plasmids were used: pcDNA3 (Invitrogen), pEPkan-S (Addgene plasmid #41017), and pCC1BAC-his3 [64,65]. pcDNA-m119-118-HA, pcDNA-m138-FLAG, and pcDNA-gB-FLAG were generated by inserting the PCR-amplified coding sequences between the BamHI and EcoRI sites of pcDNA3. The HA or FLAG tag sequences were included in the reverse primers. To generate pCC1BAC-MCMV, two 60 bp homology hooks corresponding to the first and last 60 bp of the MCMV genome were inserted between the EcoRI and BamHI restriction sites of pCC1BAC-his3. A unique ClaI restriction site was placed between the homology hooks for linearization of the vector.

### DNA preparation and next-generation sequencing

MCMV DNA was isolated from virions (vDNA) in the supernatant of MCMV-infected 10.1 fibroblasts using proteinase K digestion, phenol/chloroform/isoamyl alcohol extraction, and isopropanol precipitation, as described previously [37]. MCMV BAC DNA was purified from 200 mL of LB overnight culture using a NucleoBond Xtra Midi kit (Macherey-Nagel) following manufacturer’s instructions.

Short-read sequencing (Illumina technology) of vDNA was done at the LIV NGS facility as described previously [37]. BAC clones were analyzed by short-read (Illumina) sequencing at LIV or long-read (Oxford Nanopore) sequencing at Eurofins Genomics (Ebersberg, Germany). All sequencing reads were *de novo* assembled using SPAdes genome assembler and aligned to the respective reference genomes using Bowtie2, as previously described [37]. The complete sequences of MCMV strains v70 and K181^+^ were deposited at GenBank (accessions QB061182, QB061183). An updated sequence of the K181^Perth^ strain is available at GenBank (accession AM886412). The complete sequences of the v70, K181^+^, and K181^Perth^ BACs generated in this study were also deposited at GenBank (accessions QB061181).

DNA sequence alignments were done using BLAST [66].

### Construction of MCMV BACs by STAR cloning

STAR cloning of MCMV genomes was done as described previously for RCMV and MnatCMV genomes [37,39]. Briefly, 1.5 µg vDNA and 500 ng of the pCC1BAC-MCMV cloning vector (linearized by ClaI digestion) were combined and used for transformation of *S. cerevisiae* strain VL6-48N (ATCC MYA-3666). Transformed spheroblasts were plated on SD-SORB/-his agar plates at 31°C for 5 to 7 days. Yeast colonies containing cloned MCMV genome were identified by PCR-screening. Plasmid DNA was isolated from PCR-positive yeast colonies using a modified alkaline lysis protocol and used to transform *E. coli* strain DH10B by electroporation. Bacteria were plated on LB agar plates containing chloramphenicol and incubated overnight at 37°C. BAC DNA of individual clones was extracted, digested, and separated by electrophoresis on 0.6% agarose/0.5×TBE gels. Reconstitution of infectious MCMV from BAC clones was done by transfecting 10.1 fibroblasts with 4 µg BAC DNA using GenJet (SignaGen) transfection reagent as described previously [67].

### Virus stocks and replication kinetics

For MCMV stock preparation, 10.1 fibroblasts were grown on 15 cm dishes and infected at an MOI of 0.02. After 5 days, 200 mL of virus-containing supernatant was harvested from 10 dishes and centrifuged for 3 h at 27,000 × *g*. Virus-containing pellets were resuspended, loaded onto a 20% sucrose cushion, and centrifuged for 1.5 h at 70,000 x *g*. Virus pellets were resuspended in 200 µl PBS, aliquoted, and stored at −80°C.

MCMV stock titers and organ titers were determined by standard plaque assay on M2-10B4 cells [18,63]. MCMV replication kinetics on 10.1 fibroblasts, J774A.1 macrophages, and iBMDMs were determined as described previously [68].

### *En passant* BAC mutagenesis

MCMV BACs were modified using the *en passant* mutagenesis method, which has been described in detail [40]. Briefly, MCMV BACs were transferred to *E. coli* strain GS1783 containing a temperature-inducible λ Red recombinase system and an L-arabinose-inducible I-SceI restriction enzyme. Linear PCR products for recombination were generated by amplifying an I-SceI/kan cassette from pEPkan-S. Oligonucleotide primers (Supplemental Table 1) were designed to contain homology arms and additional sequences (e.g., tag sequences). In two consecutive steps, the I-SceI/kan cassette was first integrated into the BAC and then removed.

### Antibodies and Western blot analysis

Antibodies against the following proteins and epitopes were used: GAPDH (14C10, Cell Signaling Technology); HA (3F10; Roche); FLAG (F7425; Sigma); MCMV IE1 (CROMA101, Capri Rijeka, Croatia); MCMV gB (M55.01, Capri), MCMV E1 (CROMA103, Capri). Secondary HRP-conjugated antibody was purchased from Agilent, Dako (goat anti-mouse IgG-HRP and swine anti-rabbit IgG-HRP), or Jackson ImmunoReseach (goat anti-rat IgG-HRP).

For Western blot analysis, cell lysates were separated by SDS-PAGE and transferred onto a nitrocellulose membrane (Amersham) by semidry blotting. In some cases, cell lysates in NP40 lysis buffer (50 mM Tris pH 7.6, 150 mM NaCl, 1% Nonidet P-40, and Complete Mini protease inhibitor cocktail [Roche]) were digested with PNGase F (New England Biolabs) following the manufacturer’s protocol. Proteins of interest were detected with protein- or tag-specific primary antibodies and HRP-coupled secondary antibodies. An enhanced chemiluminescence substrate (Amersham) supplemented with 10% of Lumigen TMA-6 (Bioquote) was used, and signals were detected using a ChemiDoc chemiluminescence imager (Bio-Rad).

### Immunoprecipitation

For transfection experiments, 4×10^6^ HEK293 cells were transfected by mixing 8 µg of purified plasmid DNA with 32 µl polyethylenimine (PEI, Sigma). After incubation for 20 min at RT, the mixture was added dropwise to the cells. The medium was changed after 24 h, and the cells were harvested 48 h after transfection in NP-40 buffer on ice. For infection experiments, 1×10^6^ 10.1 fibroblasts were MCMV-infected at an MOI of 1. After 48 h, the cells were washed with PBS and harvested in NP-40 buffer on ice. Cell debris was removed by centrifugation. Lysates were pre-cleared using protein G Sepharose (Cytiva Life Sciences) or streptavidin beads (Thermo Fisher Scientific) and incubated with 4 µg total mouse IgG (Sigma) or 10 µg biotinylated Fc or Fab fragments (Rockland). Precipitation was done using 70 µl of protein G Sepharose or streptavidin beads. The samples were washed three times with IP buffer I (10 mM Tris pH 7.6, 150 mM NaCl, 0.2% NP-40, 2 mM EDTA-Na_2_), twice with IP buffer II (10 mM Tris pH 7.6, 500 mM NaCl, 0.2% NP-40, 2 mM EDTA-Na_2_), and once with IP buffer III (10 mM Tris pH 7.6). Precipitates were eluted using SDS-PAGE sample buffer (125 mM Tris pH 6.8, 4% SDS, 20% glycerol, 10% 2-mercaptoethanol, 0.002% bromophenol blue) and denatured at 95°C for 10 min. Samples containing m138-FLAG were only heated to 60°C to avoid protein aggregation.

### Flow cytometry

To measure infectivity, 10.1 fibroblasts, J774A.1 macrophages and iBMDMs were MCMV-infected at an MOI of 0.5. The cells were harvested 24 hpi, washed with PBS, and fixed for 20 min at RT in PBS/2% paraformaldehyde (PFA). The cells were permeabilized in PBS containing 0.3% Triton-X100 and 2% FCS, and treated with Fc block (TruStrain FcX, BioLegend; dilution 1:200) for 20 min at 4°C to avoid unspecific staining. After washing, the cells were incubated with anti-IE1 (CROMA101, Capri Rijeka, Croatia) for 30 min at RT. The cells were washed twice in permeabilization buffer and incubated with anti-mouse IgG coupled to AlexaFluor555 (Thermo Fisher Scientific) for 30 min at RT. The cells were washed twice in FACS buffer (PBS, 1mM EDTA, 1% FCS, 0.5% PFA) and analyzed using a LSRFortessa flow cytometer (BD Biosciences).

To measure Fc cell surface binding, 10.1 cells were infected and harvested using PBS containing 2 mM EDTA. The cells were washed once in PBS and incubated with biotinylated Fc fragment (diluted 1:100, in PBS containing 2% FCS) for 30 min at RT. They were washed twice in PBS/2% FCS and incubated with PE-labeled streptavidin (BioLegend, diluted 1:100 in PBS/2% FCS) for 25 min at 4°C. They were then washed twice and fixed with PBS/2% PFA for 15 min at RT. Finally, they were washed with FACS buffer and analyzed by flow cytometry. Data were acquired using a FACSDiva software (BD Biosciences) and analyzed using FlowJo.

### MCMV virion purification

MCMV v70 expressing tagged vFcRs were purified using a discontinuous Nycodenz (Sigma) gradient as described previously [69]. Briefly, virus was isolated from the supernatant of MCMV-infected 10.1 fibroblasts. After removal of cell debris by centrifugation at 7,500 × *g* for 15 min, virus was purified using a sorbitol cushion (20% D-sorbitol, 50 mM Tris pH 7.4, 1 mM MgCl_2_) and centrifugation at 70,000 × *g* for 1.5 h at 4°C. Virus was further purified by using a discontinuous 5-50% Nycodenz gradient. The virus-containing fraction was identified as a faint pale band at the interface between the 40% and 50% Nycodenz layers and was extracted by a syringe inserted through the side of the centrifuge tube. The particles were washed using a 20% sucrose gradient and lysed in 100 µl NP-40 buffer. Lysates were mixed with 100 µl SDS-PAGE sample buffer for Western blot analysis.

### Host FcR reporter cell assay

The assay was performed as described previously [50,51]. Briefly, 10.1 fibroblasts were seeded in 96-well plates and infected with MCMV at an MOI of 1. At 24 hpi, the cells were washed with PBS and incubated for 30 min at 37°C with a serial dilution of serum obtained from MCMV-infected or naïve BALB/c mice in RPMI supplemented with 10% FCS, 5% CO_2_. BW5147 reporter cells expressing FcγR-CD3ζ chimeric receptors were added to the cells for 18 h. Reporter cell IL-2 secretion was quantified using an α-IL-2 sandwich ELISA as described previously [50]. As a variation of this standard assay, purified virions were directly immobilized on the ELISA plate [70]. Briefly, virions (30,000 PFU per well) were used to coat ELISA microtiter plates (Thermo Fisher Scientific) in PBS for 2 h at 37°C and blocked with PBS containing 10% FCS for 1 h at 37°C, before reporter cells were added as described. For the hCD8-based FcR reporter assay, hCD8^+^ or hCD8^-^ CIM cells were infected with MCMV for 18 h, washed with PBS and incubated with serial dilutions of anti-hCD8 antibody (OKT8, Thermo Fisher Scientific) in RPMI for 30 min at 37°C.

### Ethics statement

All animal experiments were performed according to the recommendations and guidelines of the FELASA (Federation for Laboratory Animal Science Associations) and Society of Laboratory Animals (GV-SOLAS) and approved by the institutional review board and local authorities (Behörde für Gesundheit und Verbraucherschutz, Amt für Verbraucherschutz, Freie und Hansestadt Hamburg, reference number N093/2024) or the local animal welfare committee (Regierumgspräsidium of Freiburg, reference number 35-9185.81/G-14/109).

### Mouse infection experiments

For anti-MCMV serum production, BALB/c mice (Janvier Laboratories, mixed sex, 8-12 weeks old) were infected intraperitoneally with MCMV Smith strain. Blood was collected at days 42, 56 and 72 post infection. Serum was isolated by centrifugation for 10 min at 10,000 × *g* and heat-inactivated for 30 min at 56°C. Naïve serum was obtained in the same way from uninfected control mice.

For in vivo infection experiments, 6-8 week-old female BALB/c mice (Janvier Laboratories) were MCMV-infected intraperitoneally with 2×10^5^ PFU per mouse as described previously [15,18]. Organs were harvested on 3, 7, and 14 dpi, homogenized, and stored at −80°C.

### Statistical analysis

Statistical analyses were performed using GraphPad Prism versions 5. Flow cytometric Fc-binding data as well as reporter assays for measurement of host FcR activation were analyzed using one-way ANOVA followed by Bonferroni’s multiple-comparison test. Viral titers in organs after *in vivo* infection were analyzed by two-way ANOVA with Bonferroni’s multiple-comparison test of log_10_-transformed data. Adjusted *p* values < 0.05 were considered statistically significant.

## Data availability

Viral genome and BAC sequences generated in this study have been deposited at GenBank under the following accessions: QB061181, QB061182, QB061183. All other data underlying the findings reported in this study are provided in the manuscript and its supporting information files.

## Acknowledgments

We thank Edward Mocarski for providing the MCMV strains v70 and K181^Perth^, Jason Upton for MCMV K181^+^, and Zsolt Ruzsics for mouse sera. We acknowledge support from the LIV Flow Cytometry, Small Animal, and Next-Generation Sequencing facilities. The LIV is supported by the Free and Hanseatic City of Hamburg and the Federal Ministry of Health.

## Author contributions

Conceptualization: LS, PK, HH, and WB; investigation: LS, MM, and EO; formal analysis: AJD; resources: KE, BC, AJR, PK, and HH; supervision: FRS, HH, and WB; writing - original draft: LS and WB; writing - review & editing: all authors.

## Supplementary data

**S1 Table**. **List of oligonucleotides and gBlocks used in this study.** Underlined sequences indicate restriction enzyme cleavage sites.

**S1 Fig. Comparison of viral replication and infectivity of MCMV strains.** (**A**) J774A.1 macrophages were infected with MCMV strains v70, K181^+^, or K181^Perth^ (MOI=0.5). (**B**) iBMDMs were infected as in (A). (**C**) 10.1 fibroblasts were infected with MCMV strains v70, K181^+^ and K181^Perth^ (MOI=0.02). Viral titers in the supernatant were determined using the TCID_50_ method. Mean ±SEM of three biological replicates is shown. DL, detection limit. (**D-F**) J774A.1 macrophages, iBMDMs, and 10.1 fibroblasts, respectively, were infected with MCMV v70, K181^+^, or K181^Perth^ (MOI=1.5) for 24 h. The cells were fixed with PFA, permeabilized and stained with an anti-IE1 antibody. Infectivity was measured by flow cytometry. The results of three independent experiments are shown.

**S2 Fig. Quantification of virion proteins by ELISA.** Adjusted amounts of purified virions of v70 WT and various mutants were used to coat ELISA plates. Coated plates were incubated with anti-MCMV antiserum from BALB/c mice, followed by incubation with HRP-coupled anti-mouse-IgG. Viral proteins per well were quantified by ELISA. Uncoated wells served as negative control (blank). Mean ±SEM from three independent experiments are shown.

**S3 Fig. CD8 expression by transduced CIM cells.** CIM hCD8+ and non-transduced parental CIM hCD8-cells were stained with APC-labeled antibody against hCD8 or mock-stained. APC staining was analyzed by flow cytometry.

